# Oxidative stress delays development and alters gene function in the agricultural pest moth, *Helicoverpa armigera*

**DOI:** 10.1101/2020.01.14.906958

**Authors:** Nonthakorn (Beatrice) Apirajkamol, Bill James, Tom K Walsh, Angela McGaughran

## Abstract

Stress is a widespread phenomenon that all organisms must endure. Common in nature is oxidative stress, which can interrupt cell homeostasis to cause cell damage and may be derived from respiration or from environmental exposure thought diet. As a result of the routine exposure from respiration, many organisms can mitigate the effects of oxidative stress, but less is known about responses to oxidative stress from other sources. *Helicoverpa armigera* is a major agricultural pest moth that causes significant damage to crops worldwide. Here, we examined the effects of oxidative stress on *H. armigera* by chronically exposing individuals to paraquat - a free radical producer - and measuring changes in development (weight, developmental rate, lifespan), and gene expression.

We found that oxidative stress strongly affected development in *H. armigera*, with stressed samples spending more time as caterpillars than control samples (>24 vs. ∼15 days, respectively) and living longer overall. We found 1,618 up- and 761 down-regulated genes, respectively, in stressed vs. control samples. In the up-regulated gene set were genes associated with cell senescence and apoptosis and an over-representation of biological processes related to cuticle and chitin development, glycine metabolism, and oxidation-reduction.

Oxidative stress clearly impacts physiology and biochemistry in *H. armigera* and the interesting finding of an extended lifespan in stressed individuals could demonstrate hormesis, the process whereby toxic compounds can actually be beneficial at low doses. Collectively, our findings provide new insights into genomic responses to oxidative stress in invertebrates.

## Introduction

Stress is encountered by all living beings with its various manifestations producing different responses, from small-scale molecular changes to large-scale shifts in development and lifespan. In a variety of species, stress has been shown to strongly affect fitness, resulting in changes in organismal behaviour, developmental rate, physiology, and mortality (Adamo, 2012; Trakimas et al., 2019). For example, McCormick et al. (1998) reported that physical stress (chasing, crowding, and draining) in Atlantic salmon resulted in lower growth rates and body weight. In addition, stress has been shown to shorten lifespan in some species (e.g., humans, Shalev et al., 2013; cane toads, Jessop et al., 2013), but to extend lifespan in others (e.g., *Drosophila melanogaster*, Hercus et al., 2003; Jessop et al., 2013; *Caenorhabditis elegans*, Lithgow & Walker, 2002). As well as impacting development, stressful environments can alter cell-signalling pathways, resulting in changes in gene expression, metabolism, cell cycles, protein homeostasis, and enzyme activity (Rampon et al., 2000; Richter et al., 2010; Weake & Workman, 2010; Nadal et al., 2011).

An important component of the overall stress response is oxidative stress, which is a toxic by-product of aerobic metabolism (Lushchak, 2014). Oxidative stress occurs when oxygen becomes excited and hyperactive i.e., reactive oxygen species - ROS (Kregel & Zhang, 2007), reacting with other molecules (Halliwell & Gutteridge, 1984; Imlay, 2003) to increase free radical (e.g., hydroxyl radicals, superoxide anions, and hydrogen peroxide) production (Finkel & Holbrook, 2000). As a consequence, the balance between antioxidant production and ROS removal can be disrupted, ultimately causing damage to cellular components such as DNA, enzymes, and cell membranes (Betteridge, 2000; Mittler, 2002; Blumberg, 2004). The majority of aerobic organisms have developed sophisticated methods to relieve the effects of oxidative stress (Krishnan et al., 2007), however, little is known about this process in pests.

*Helicoverpa armigera (*Lepidoptera: Noctuidae), is a major agricultural pest moth. Its’ larvae cause damage by consuming the reproductive parts of plants and *H. armigera* can feed on >300 different host species, collectively causing the agricultural industry losses of ∼$USD 5 billion annually (Pearce et al., 2017). *H. armigera* has a wide environmental tolerance range, a high fecundity, and is able to migrate over very large (>1000 km) distances (Feng et al., 2004). These factors together enable *H. armigera* to occupy a worldwide distribution encompassing Asia, Australia, Africa, Europe, and more recently, parts of South America (Czepak et al., 2013; Kriticos et al., 2015).

General information about stress responses in *H. armigera* is scarce. However, various measures of development have been shown to respond to stress in this species. For example, both weight and developmental rate are associated with the type of host plant on which individuals are raised. In particular, caterpillars reared on less favourable host plants (e.g. *Arabidopsis*, tobacco, and tomato) take longer to reach certain developmental stages and also have a lower body weight (Pearce et al., 2017). Yet, in other research, different types of stress have been shown to affect *H. armigera* differently. For example, higher temperatures and increased predator stress have been shown to hasten development (Xiong et al., 2015; Noor-Ul-Ane et al., 2018), whereas low temperatures, and poor diet apparently elongate developmental periods in *H. armigera* (Xiong et al., 2015; Pearce et al., 2017; Noor-Ul-Ane et al., 2018). With respect to oxidative stress, acute exposure has been shown to extend lifespan in *H. armigera* (Zhang et al., 2017) and, when exposed to ultraviolet (UV) radiation (a common environmental stress that increases levels of oxidation), *H. armigera* shows up-regulation of antioxidant genes, but only at certain UV doses (Wang et al., 2012).

Only few studies have targeted the effects of oxidative stress on *H. armigera* and none have looked at chronic exposure. This is unfortunate because oxidative stress is often associated with acute exposure to pesticides (Abdollahi et al., 2004), and is faced chronically by *H. armigera* in the wild via natural plant defence mechanisms (many plant species are able to intensify levels of oxidative stress to defend themselves against herbivore and virus attacks; e.g. Aucoin et al., 1991). In this study, we examine the effects of oxidative stress on *H. armigera* using the oxidative producer, paraquat.

Paraquat (*N,N*′-dimethyl-4,4′-bipyridinium dichloride) is an organic herbicide that kills a wide range of pests by generating superoxide anions (Shadnia et al., 2018). Despite being banned in several countries, paraquat is widely available throughout the world due to its efficiency and low cost (Kim et al., 2017), and there is evidence that it has a strong impact on development and gene expression in some insects. For example, in *D. melanogaster*, paraquat has been shown to increase mortality, reduce climbing ability, and result in up-regulation of several antioxidant genes (Krůček et al., 2015). In *H. armigera*, paraquat injected into pupae has been shown to extend diapause by affecting the insulin-signalling pathway (Zhang et al., 2017), but no work to date has focused on stress effects of paraquat in other developmental stages of *H. armigera*. Thus, we examine development and gene expression in *H. armigera* following chronic oxidative stress exposure. Our results provide fundamental information about genomic responses to stress in invertebrates and new insights into how changes in development may shape population dynamics, ecosystem consequences, and evolutionary adaptation in this species.

## Methods

### Insect rearing

All experiments used a lab colony of *H. armigera conferta*, which is widespread across Australia and New Zealand (Anderson et al., 2016). The colony was originally established from cotton fields in the Namoi Valley, northern NSW Australia and has been reared in lab conditions at the Commonwealth Scientific and Industrial Research Organisation (CSIRO) in Canberra since the mid-1980s.

To initiate experiments, a large number of two-day old fertilised eggs were collected from 40 healthy moths and disinfected by soaking in a 0.002% bleach solution (0.0008 mv^-1^ of chlorine) for 10 min and then washing with tap water. Eggs were air-dried and placed in a plastic bag to allow hatching. Subsequently, 1st-instar caterpillars were transferred to 32-well plates for rearing until pupation under optimum conditions (25±1°C, 50±10% relative humidity, and light day:night 14:10 to imitate natural light), with a solid artificial diet (see below), which was changed every week to prevent stress from insufficient food. Once pupated, the sex of individuals was determined under the microscope. Two pupae of the same sex were then placed into individual containers (separated by paper to prevent interaction) until they died. Moths were fed a honey solution as per normal rearing protocols (see below), which was checked every two days and refilled if needed.

The semi-solid artificial diet was prepared for caterpillar rearing, using an in-house protocol. First, 130 g of soy flour and 700 mL of filtered water were blended with a stick blender until combined. The mixture was then heated in the microwave until it boiled (4-5 min). Next, 22 g of agar (GELITA, A-181017), 1.7 g of sorbic acid (Sigma-Aldrich, S1626), and 700 mL of filtered water were combined and gently mixed with a spatula. This mixture was also heated in the microwave to boiling (4-5 min). Both mixtures were then separately stirred and reheated before being combined together by blending with a stick blender. Additional dry ingredients, including 60 g of wheat germ, 53 g of brewers dry yeast, 3.3 g of L-ascorbic acid (heat sensitive, so after cooling to 60°C; Sigma-Aldrich, A4403), and 3.3 g of nipagin (Methylparaben, Sigma-Aldrich, 79721), were added to the final mixture, along with 5 mL of vegetable oil. Filtered water was added to bring the total volume to 1600 mL and the mixture was blended until well-combined. This diet was poured into 32-well plates (approximately 5 mL per well) and left at room temperature for at least an hour to dry, cool down, and set. In all cases, the diet was immediately used or stored at 4°C for a maximum of two days.

After emerging, *H. armigera* moths were fed on a honey solution. To prepare the honey solution, 40 g of white sugar, 40 g of honey, and 1 g of sorbic acid were weighed into a 1 L bottle. Around 300-400 mL of filtered water was added, then the mixture was heated in the microwave for 2 min on high heat and shaken to dissolve the sugar. Filtered water was added to bring the final volume to 1 L; the mixture was then shaken and, with a loosened lid, placed in an autoclave for 15 min at 121°C to sterilise. After cooling to 60°C, 2 g of ascorbic acid was added to the mix, which was then shaken and stored at 4°C.

### Experimental design

To determine the optimal ‘stress’ conditions to expose *H. armigera* to, a range-finding experiment was performed, where individuals were exposed to a number of different concentrations of paraquat (Sigma-Aldrich 36541) through their diet (i.e., added into the solid artificial diet before it set or directly into the honey solution). A response was observed (reduction in weight and body size) in samples exposed to >0.25 mM of paraquat, while individuals exposed to 0.5 mM had an overall mortality exceeding 50%. Therefore, a final selection of paraquat concentrations 0.3 mM and 0.4 mM was made to create moderately stressful conditions (i.e. based on weight/mortality/developmental rate). Individuals were split into three groups, corresponding to control (normal diet as outlined above), and stressed (0.3 mM or 0.4 mM paraquat) and examined for developmental phenotypes and gene expression.

### Developmental phenotypes

In each treatment group and control, individuals were randomly selected for assessment of weight (n = 10), developmental stage (n = 32), and mortality (n = 32). Each of these phenotypes was recorded for the same individuals every two days across three replications (e.g., total n = 96 for developmental stage and mortality) until all samples had died. In addition, the amount of time spent in each developmental stage (caterpillar, pupa, and moth), and the overall lifespan (time from hatching to death) was recorded for 40 males and 40 females for each treatment group and control.

### Statistical analysis

Developmental measures were statistically analysed via SPSS ver. 22 (IBM Corp, 2013) to determine whether there were any differences between control and stressed groups. Mean and standard deviations (SD) were calculated for all measures and outliers (individuals with values greater/lesser than mean ∓ 2 SD) were removed. One-way ANOVA with Tukey’s HSD (Honestly Significant Difference) was used to assess differences between groups at a 95% confidence interval. The Tukey HSD tests were used in order to reduce false positives from family-wise error rates (due to the number of treatments), and type II errors from a large number of standard deviations (Kaufmann & Schering, 2014). In this analysis, samples that are statistically similar (in terms of mean and variance for the trait being measured) group together into homogeneous subsets. Thus, if samples are categorised into different groups (referred to as ‘a’, ‘b’, ‘c’, etc.; see Results), they are considered significantly different from each other. Data visualisations were performed using the ggplot2 ver. 3.2.1 (Wickham, 2016) package in R ver. 3.6.1 (R Core Team, 2019).

### Gene expression analysis

#### Sample collection and RNA isolation

A total of 72 samples were used for gene expression analysis, corresponding to three groups (controls, 0.3 mM and 0.4 mM paraquat), with four replicates per group, and each replicate consisting of a pool of six individuals. Care was taken to select samples of similar body size for each pool. Each sample was collected at 4th-instar, snap frozen with liquid nitrogen, and stored at −80°C. Whole caterpillars were then homogenised in TissueLyser II solution (Qiagen), with 12 × 2 mm ceria-stabilised zirconium oxide ceramic beads (ZROB20-RNA, Next Advance) in 500 µl of 90% ethanol at 30 ls^-1^ for 3 min – where the beads, ethanol and TissueLyser sample racks were pre-chilled to −80°C. Samples were then re-chilled on dry ice for 1 min and the homogenisation process repeated twice. Subsequently, 20-120 µl of combined homogenates from six individuals (totalling 10 mg of tissue per individual) were aliquoted into a pool and thoroughly mixed. Into each pool, 1200 µl of Lysis Buffer (PureLink™ RNA Mini Kit) was added, along with 70% ethanol to bring the total to 1600 µl (achieving a final concentration of 40% ethanol, 50% lysis buffer, and 10% tissues). RNA isolation was conducted following the PureLink™ RNA Mini Kit protocol. The RNA pellet was re-suspended with 70 µl RNase free water and quantity/quality checked by Nanodrop (ThermoFisher) and MultiNA (Shimadzu).

#### RNA library preparation and sequencing

Library preparation was conducted using an in-house method adapted from Langevin et al. (2013). mRNA enrichment was first conducted in order to reduce the amount of ribosomal RNA (rRNA) in the total RNA sample. Because there are no kits available specifically for rRNA depletion in *H. armigera*, oligo d(T) capture methods were used to enrich the concentration of poly-adenylated mRNA. Magnetic beads were prepared by first removing the liquid residue from beads by placing tubes on a magnet to discharge the supernatant. Subsequently, beads were washed with 50 µl of binding buffer, re-suspended, and the buffer discharged. Two washes were performed in total. Two rounds of mRNA enrichment were then performed. In the first round, RNA was denatured by incubating 50 µl of the total RNA at 65°C for 2 min then immediately chilling on ice. 15 µl of Oligo d(T)25 Magnetic Beads (S1419S, New England Biolabs), were combined with 50 µl of binding buffer (20 mM Tris-HCl pH 7.5, 1 M LiCl, 2 mM EDTA) and added to 50 µl of each denatured RNA sample in a V-bottom assay plate (P-96-450V-C, ThermoFisher). Samples were incubated at room temperature for 15 min on a Titramax plate shaker (Heidolph) at 1200 RPM to allow the mRNA to hybridise to the beads. A covered magnetic separator (i.e., a neodymium 50 (N50) magnet in a 3D printed case) was inserted and the plate incubated for a further 2 min. Beads were then washed with 120 µl of washing buffer (10 mM Tris-HCl pH 7.5, 150 mM LiCl, 2 mM EDTA), with the washing process repeated twice. mRNA was eluted by submerging the washed magnetic separator with the beads in 50 μl of Elution Buffer (10 mM Tris-HCl pH 7.5), then the magnetic separator was removed and the plate was incubated at 80°C for 2 min to re-suspend the beads and mRNA. Finally, the magnetic separator was then inserted to capture and remove the beads. For the second round of mRNA enrichment, 50 µl of binding buffer was added to eluted mRNA, then the washed oligo d(T)25 magnetic beads from round one (following re-suspension in nuclease-free water at least two times) were transferred into the wells. Samples were incubated on a Titramax plate shaker at 1200 RPM for 15 min and the magnetic beads were captured with the separator. The beads were washed by transferring them with the separator to 20 μl of Washing Buffer 2, repeated twice. Subsequently, mRNA was eluted by transferring the washed beads with separator to a new well with 8 μl of Elution Buffer and the magnetic separator was removed. Samples were finally incubated at 80°C for 2 min, beads were removed by capturing them with the magnetic separator, and the purified mRNA was stored at −80°C.

Fragmentation was conducted in order to ensure the desired insert sizes for Illumina sequencing, by combining heat and Mg^2+^ ions with the mRNA. Fragmented mRNA was then synthesised to first strand cDNA using a reverse transcription process. 3 μl of purified mRNA was combined with 1 μl of 5× SMARTscribe RTase buffer (TakaraBio) and 1 μl of 1 μM RT-Hex primer and incubated at 85°C for 5 min. Samples were immediately transferred onto ice for 2 min and then let rest at room temperature for at least 10 min. Finally, 5 μl RT master mix (1 μl of 5x SMARTscribe RTase buffer, 1 μl of 10 mM dNTPs, 0.5 μl of 100 mM DDT, 0.25 μl of RiboLock RNase inhibitor, 1 μl of 10 μM Bio_TS_RNA primer, 0.5 μl of SMARTscribe RTase, and 0.75 μl of nuclease-free water) was added to the fragmented mRNA and mixed thoroughly. Samples were incubated in a thermal cycler for reverse transcription and template switching reactions to occur, under the following cycling conditions: 25°C for 30 min, 42°C for 90 min, 72°C for 10 min, with a heated lid temperature of 45°C throughout and a starting block temperature of 25°C.

First strand cDNA samples were purified using solid phase reversible immobilisation (SPRI) paramagnetic MagNA magnetic bead (Rohland & Reich, 2012). The completed reverse transcriptase reactions were transferred to V-bottom assay plates, then 9.5 μl of MagNA beads were added. Samples were thoroughly mixed and incubated at room temperature for 8 min. The MagNA beads were recaptured by inserting a magnetic separator, then incubated at room temperature for at least 2 min. The magnetic separator and beads were then transferred into 200 µl of 80% ethanol for washing. Beads were washed twice and then allowed to air dry for at least 3 min. cDNA was re-suspended by transferring the washed magnetic separator with the beads into 10 µl of nuclease-free water. The separator was then removed and the sample incubated for 5 min. Beads were again recaptured by inserting the magnetic separator and then discarding it. Finally, the cDNA MagNA magnetic clean-up process was repeated in order to ensure complete depletion of short, empty constructs, this time with a final elution volume of 20 µl. Purified cDNA was processed immediately or stored at −20°C.

The optimal cycle number for the barcoding of each individual sample was determined by qPCR. The purified first strand cDNA was diluted by aliquoting 1 μl of cDNA into 15 μl of nuclease-free water. A qPCR reaction was set up in a 10 μl reaction volume (5 μl of Bio-Rad SsoFast EvaGreen Supermix, 1 μl of 2.5 μM TS_qPCR and 2.5 μM RT_Hex_qPCR primer mix, 4 μl of diluted cDNA template), and then cycled on a Bio-Rad CFX96 thermal cycler with the following conditions: 95°C for 45 s, followed by 35 cycles of 95°C for 5 s and 60°C for 30 s. The optimal cycle number of the undiluted 1st strand of cDNA was calculated based on the quantification cycle (Cq) number of the diluted samples.

Each sample was then barcoded with a unique pair of indexed primers. A PCR master mix was prepared for a 7 μl per-sample reaction volume (4 μl of 5x Phusion buffer, 0.4 μl of 10 mM dNTPs, 2.4 μl of nuclease-free water, and 0.2 μl of Phusion polymerase), then mixed with 8 μl of purified (undiluted) first strand cDNA. 2.5 μl each of forward and reverse barcode primers was thoroughly mixed into each individual sample and PCR was performed according to the predetermined optimum cycle number according to the following: 1 cycle of 98°C for 10 s (with the block pre-heated to 80°C), *x* cycles of 98°C for 5 s, 58°C for 10 s, 72°C for 20 s, then 1 cycle of 72°C for 5 min, where *x* is the optimal cycle number specific to each sample.

After the barcoding process, cDNA was cleaned up with MagNA magnetic beads similarly to the first strand cDNA clean-up, but with 17 μl of beads instead of 9.5 μl, the replacement of 80% ethanol with 70% ice-cold ethanol, and the elution of DNA with 20 μl of 10 mM Tris pH 8.0 instead of nuclease-free water.

Barcoded samples were serially diluted to a 1:10,000 dilution and sample quantity was determined using Library Quant Master Mix for Illumina (NEBNext^®^ E7630). qPCR reactions were set up in duplicate on the diluted template according to the kit instructions, except all volumes were halved. Samples were then pooled according to the qPCR quantifications to achieve equimolarity. The pooled, equimolar, cDNA library was sequenced using a custom read 1 primer at the Biomolecular Resource Facility (BRF) at Australian National University on a NovaSeq6000 SP machine (2 × 50 bp paired end sequencing). All primers used during this protocol are documented in the Supporting Information (Table S1).

#### Gene expression analysis

Raw sequence reads were checked for quality using FastQC (Andrews, 2010; freely available at: https://www.bioinformatics.babraham.ac.uk/projects/download.html#fastqc) and then mapped to the *H. armigera* reference genome using STAR ver. 2.7.2b (Dobin et al., 2013). STAR was also used to produce a table of gene counts using default settings.

Differential gene expression (DGE) analysis was performed in R using various packages, including edgeR ver. 3.26.8 (Robinson et al., 2010), limma ver. 3.40.6 (Smyth, 2005), and ggplot2. First, genes that were unexpressed or not expressed at biologically meaningful levels were filtered in order to reduce mean-variance from low count data in further analysis. For a gene to be retained, it needed to be counted at least ten times and present across at least two replicates. Gene expression distributions were then normalised with calcNormFactors from edgeR to ensure that differences in sequencing depth between replicates did not skew results. A multi-dimensional scaling (MDS) analysis was performed using the function plotMDS from limma to visualise differences and similarities between samples in the top 1,000 most highly-expressed genes. Finally, DGE analysis was performed using the voom workflow from limma and a list of the DE (up- and down-regulated) genes was generated.

#### Gene ontology analysis

Gene Ontology (GO) analysis was performed to indicate which GO terms were over- or under-represented in the table of DE genes. This analysis was conducted in R, using the doSNOW ver. 1.0.18 (available at: https://cran.r-project.org/web/packages/doSNOW/index.html) and foreach ver. 1.4.7 (available at: https://cran.r-project.org/web/packages/foreach/index.html) packages. A custom script developed by Dr. Darren Wong was used to match the gene names of DE genes against a table of characterised GO terms using a mapping file obtained from Pearce et al. (2017).

## Results

### Developmental phenotypes

#### Weight

The mean weight profile of control and stressed samples is shown in Figure 1. All samples increased in mean weight to ∼0.4-0.5 g as growing caterpillars, then declined towards a final lower mean weight (∼0.15 g) that was relatively consistent among treatment and control groups (Fig. 1A). Though the general pattern was similar for all samples/treatments, there were some differences. Firstly, the mean weight of control samples more rapidly increased at the beginning of the experiment. Secondly, the timing of peaks and declines differed between the three groups, for example control samples peaked in mean weight at day 12, while paraquat 0.3 mM and 0.4 mM peaked later (days 22, and 32, respectively; Fig. 1A). These differences were significant between the control and stressed groups - control samples had the highest mean weight from day four to day 16 (F_2,27_=47.613, F_2,26_=76.771, F_2,22_=497.884, F_2,23_=69.012 for days 4, 8, 12, and 16, respectively; *P*<0.001 for all) and from days 32 to 36 (day 32: F_2,17_=34.874, day 36: F_2,17_=14.297; *P*<0.001 for both) - but not between the two paraquat stressed groups, except for on day 20, when the 0.3 mM stressed samples were, on average, significantly heavier than their 0.4 mM counterparts (F_2,23_=7.387; *P*<0.001; Supporting Information Fig. S1). To account for differences in development, we also compared mean weight at the same developmental stage, finding no significant differences across treatment groups (F_2,20_=0.49, *P*=0.952; F_2,19_=1.064, *P*=0.365; F_2,15_=3.192, *P*=0.070 for caterpillar, pupa, and moth stages, respectively; Fig. 1B).

**Figure 1.**
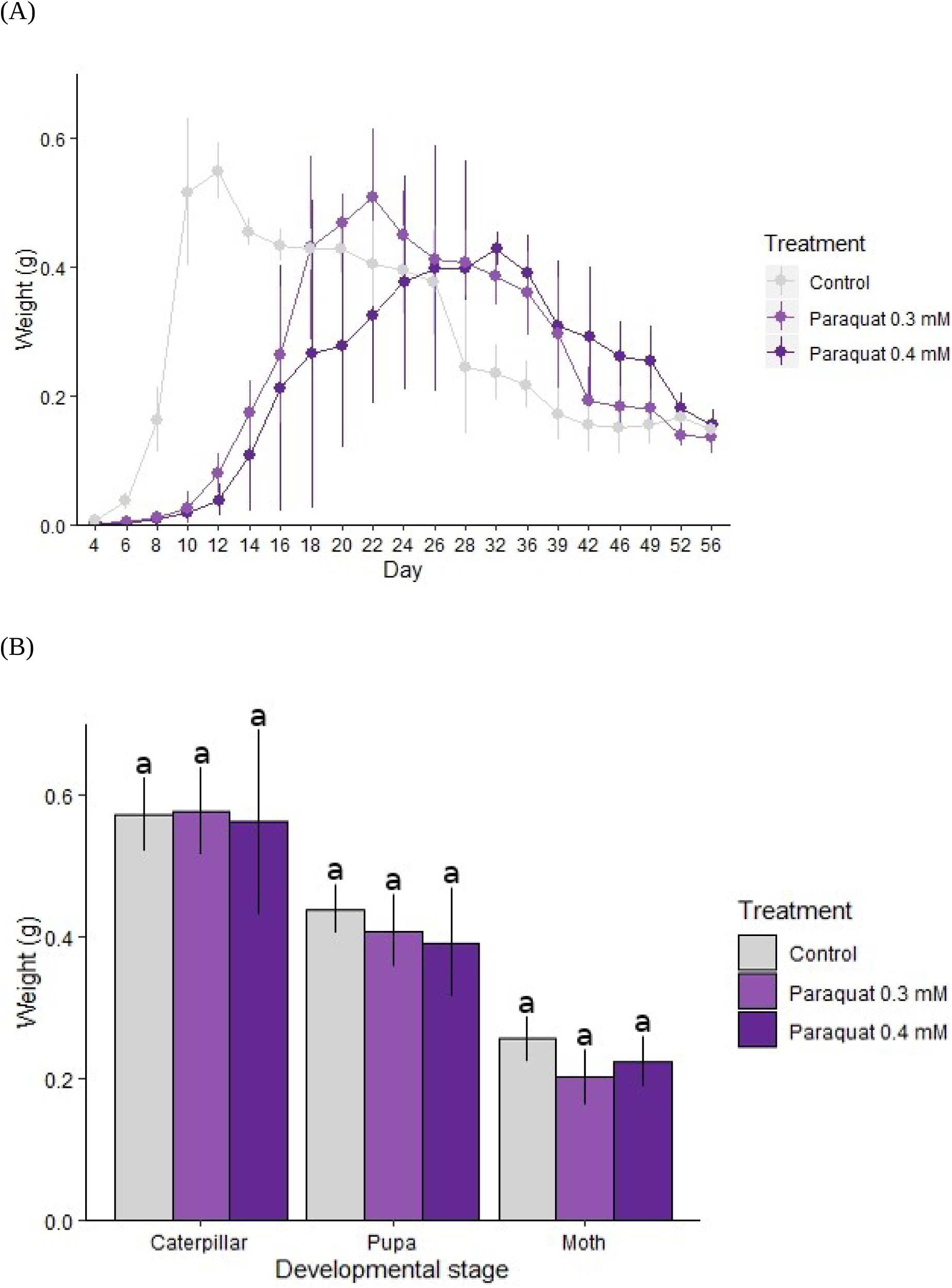
Mean weight following stress exposure in *Helicoverpa armigera*. Individual *H. armigera* were reared on an artificial diet mixed with 0.3 mM or 0.4 mM paraquat. Ten randomly-selected individuals were weighed every two days as caterpillars and twice a week as moths. Results are presented for stressed and control samples: (A) mean weight every two days from day four, and (B) at three different developmental stages. Differences between treatment and control groups in (B) were not found to be statistically significant (i.e., all fall into a single homogeneous group). In both graphs, error bars indicate standard deviation and colours represent control or treated samples according to the provided key.

#### Developmental rate

Developmental rate was measured as the percentage of samples presenting as a given developmental stage on the day of measurement. Based on this metric, paraquat-stressed samples had a significantly slower developmental rate compared to the controls (Fig. 2). For example, by day eight, more than 95% of control samples had reached 4th-instar, while the majority of stressed samples were only at either 2nd or 3rd-instar (F_2,6_=3274.973, *P*<0.001; Fig. 2A). There was a similar trend from day 12 to day 24 – control samples had consistently reached a later developmental stage than stressed samples (day 12: F_2,6_=5410.838, day 16: F_2,6_=5747.006, day 20: F_2,6_=964.347, day 24: F_2,6_=51.028; *P*<0.001 for all; Fig. 2B,C, Supporting Information Fig. S2). As for weight, the developmental rate of stressed samples was not significantly different between 0.3 mM and 0.4 mM paraquat on most measurement days (Fig. 2A, Supporting Information Fig. S2).

**Figure 2.**
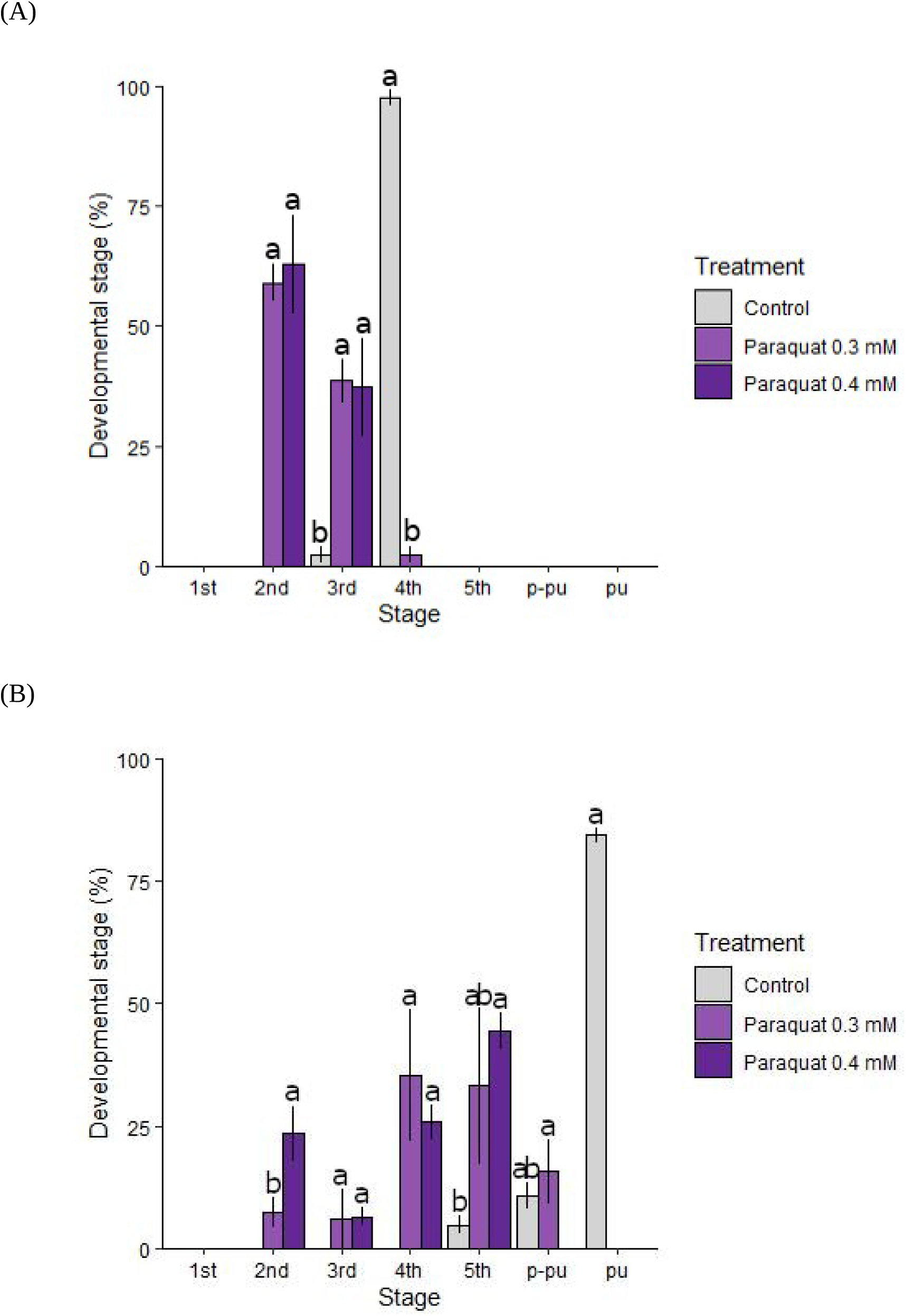
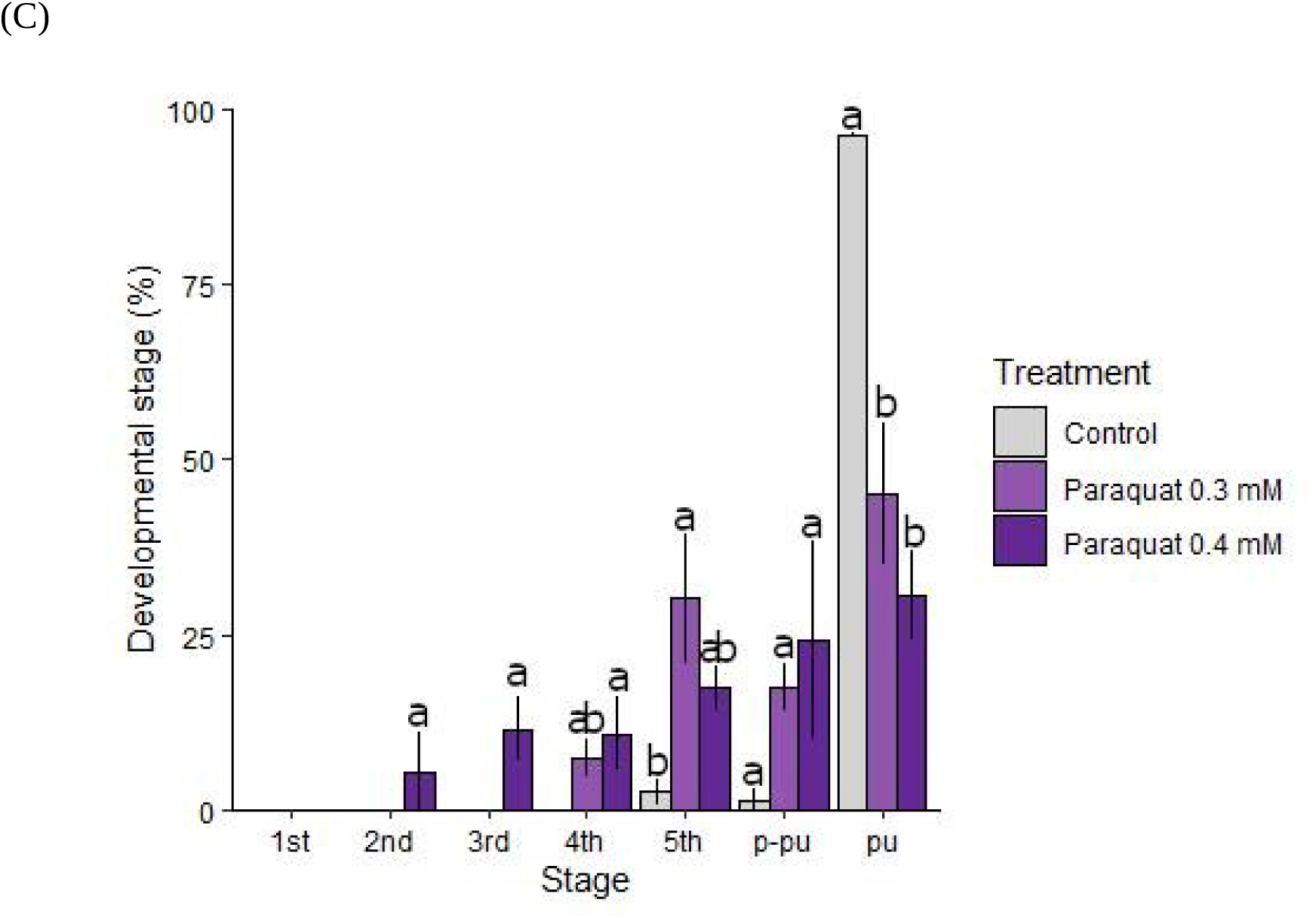
Developmental stage progression following stress exposure in *Helicoverpa armigera*. Individual *H. armigera* were reared on an artificial diet mixed with 0.3 mM or 0.4 mM paraquat and developmental stage was recorded every two days for control and stressed samples from the day that individuals hatched until death. Results are presented as the percentage of individuals representing each developmental stage (1st - 5th corresponding to instars, p-pu=pre-putation; pu=pupae) at: (A) day eight; (B) day 16; (C) day 24. Significant differences among treatment and control groups are indicated by non-overlapping characters (‘a’, ‘b’), error bars indicate standard deviation, and colours represent control or treated groups according to the provided key.

Interestingly, Figure 2 shows that stressed samples not only reached developmental stages at a slower rate, but also had higher variation in the percentage of samples present at each stage, compared to the controls. On any given day, the majority (>85%) of control samples were at the same developmental stage as each other, while stressed samples were not. For example, on day 24, ∼96% of the control samples had pupated, while the stressed samples were represented in all developmental stages except 1st-instar (Fig. 2C).

#### Lifespan

Differences in developmental rate translated into differences in average time spent in each developmental stage and in overall average lifespan across the treatment groups and controls. The mean time spent as a caterpillar (from the day of hatching to pupation) for males and females within the same treatment group was not significantly different, however the control group took significantly less time (∼15 days) to pupate than both stressed groups (∼25 and ∼28 days for 0.3 mM and 0.4 mM, respectively; F_5,276_=144.966, *P*<0.001; Fig. 3A). In contrast, there were no significant differences in pupation period among the treatment groups, but the pupation period of male samples was ∼2 days longer on average than that of females for the control group (F_5,260_=5.280, *P*<0.001; Fig. 3B). Finally, mean time spent as moths across all three groups was not significantly different (F_5,236_=0.761, *P*=0.578; Fig. 3C).

**Figure 3.**
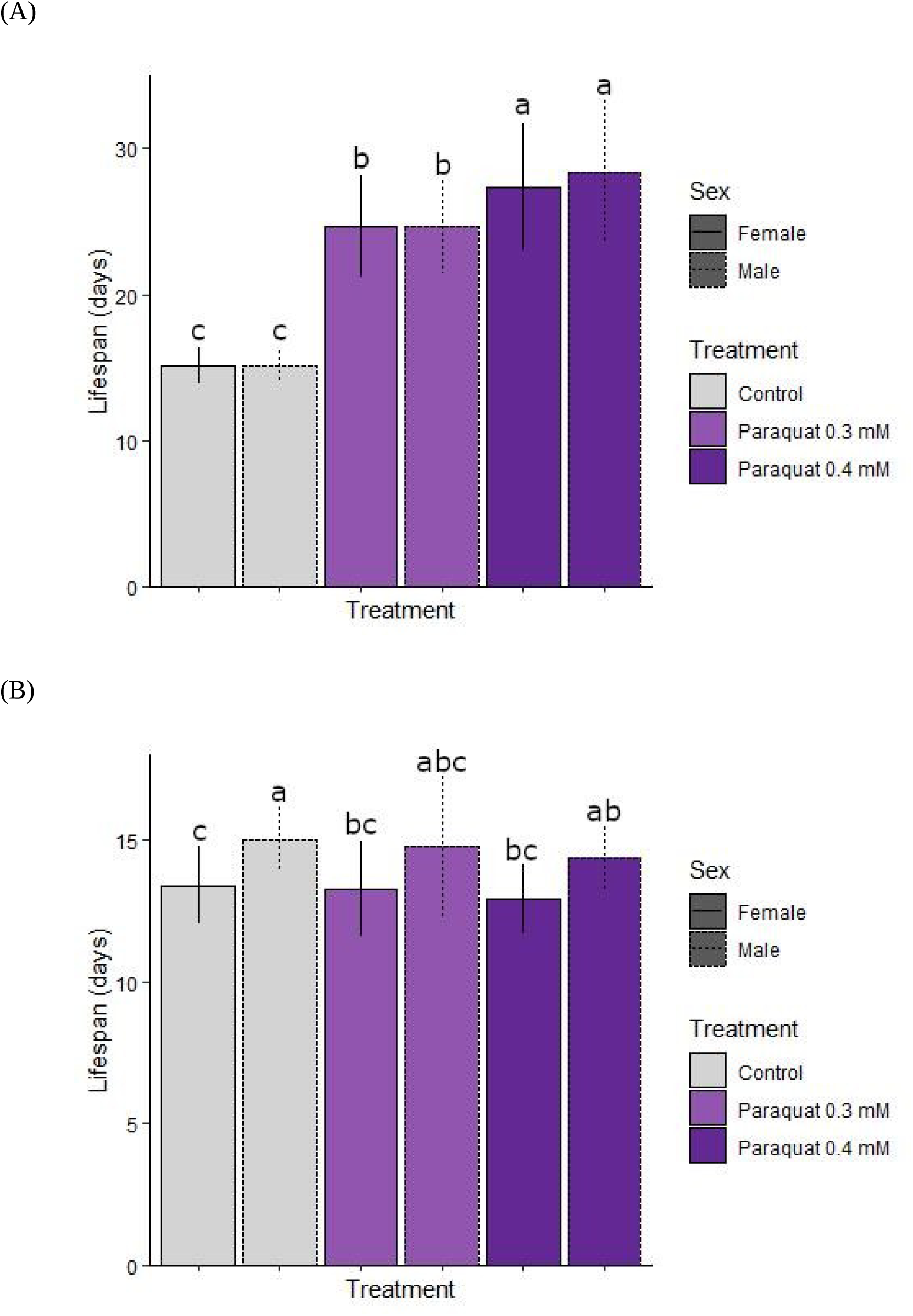
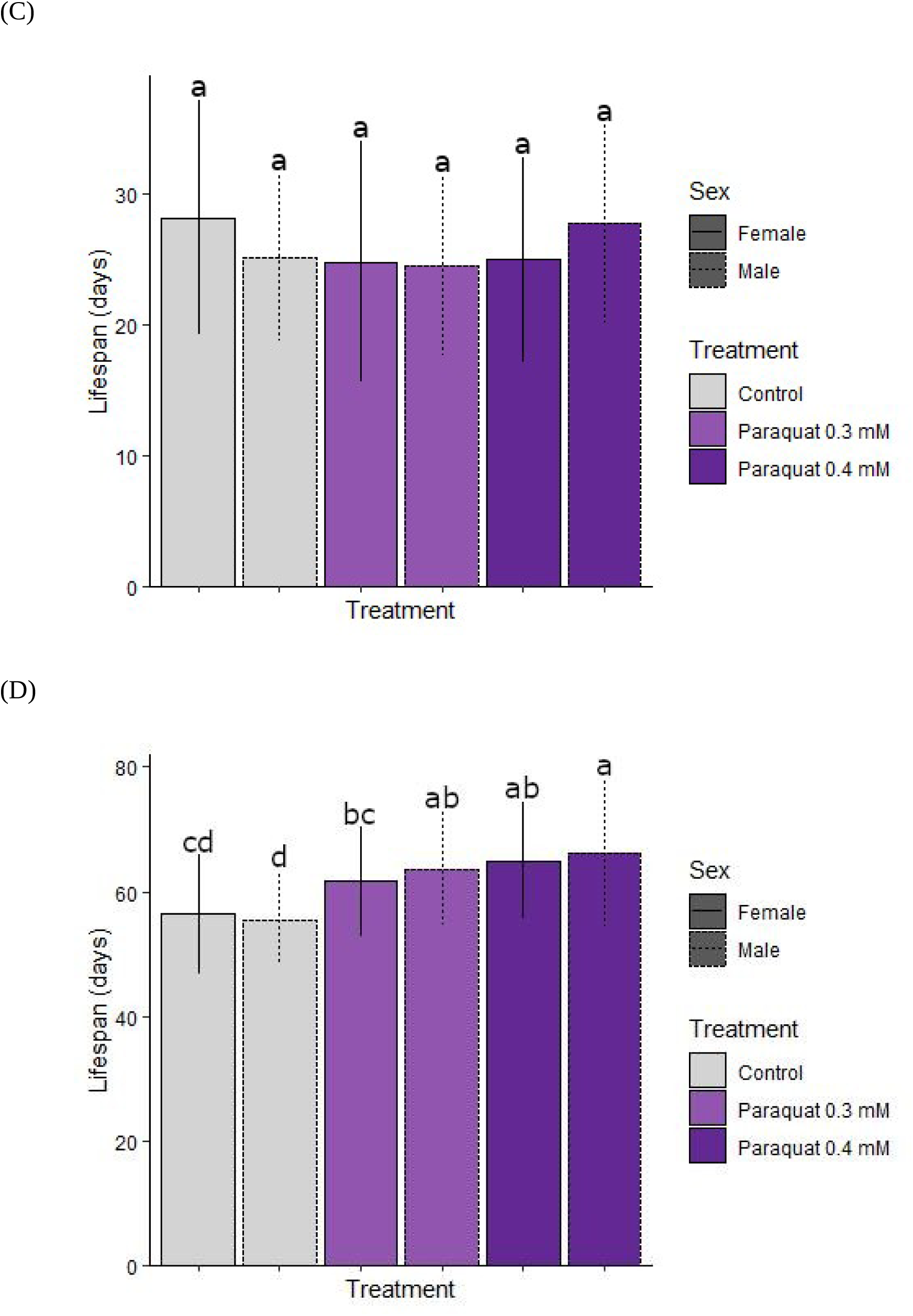
Lifespan following stress exposure in *Helicoverpa armigera*. Individual *H. armigera* were reared on an artificial diet mixed with 0.3 mM or 0.4 mM paraquat and time spent in each developmental period was recorded, as was overall lifespan (time from hatching to death). Results are presented as mean number of days spent as: (A) caterpillars; (B) pupae; and (C) moths. Overall lifespan is presented in (D). Significant differences among treatment and control groups are indicated by non-overlapping characters (‘a’, ‘b’, ‘c’, ‘d’) and error bars indicate standard deviation. Colours represent control or treated groups, and solid and dashed lines indicate females (F), and males (M), respectively, according to the provided key. Note the different *y-*axis scales across the four panels.

Overall lifespan refers to the time from hatching to death and is shown in Figure 3D, where the general pattern is an extended average lifespan in stressed samples relative to controls (F_5,234_=16.748, *P*<0.001). Specifically, the male paraquat-stressed group (0.3 mM and 0.4 mM) had a significantly longer (∼8-9 days) mean lifespan than the male and female controls and were not significantly different from each other (Fig. 3D). For females, the paraquat 0.4 mM group had a longer overall mean lifespan than the control group, but the paraquat 0.3 mM group was not significantly different to either the control or paraquat 0.4mM groups (Fig. 3D).

#### Mortality

The percent mortality profile of control and stressed samples is presented in Figure 4. All treatment groups had an individual two-daily mortality percentage of around 4-5.5% at the early stages of the experiment and this declined to <3% after day six for control and paraquat 0.3 mM groups. However, in the paraquat 0.4 mM group, two-daily mortality rates reached 3-5% for days 14, 16, 20, 24, and 26 (Fig. 4A). Overall mortality as a caterpillar reached ∼16-26% and was not significantly different across treatments (F_2,6_=1.264, *P*=0.348; Fig. 4B).

**Figure 4.**
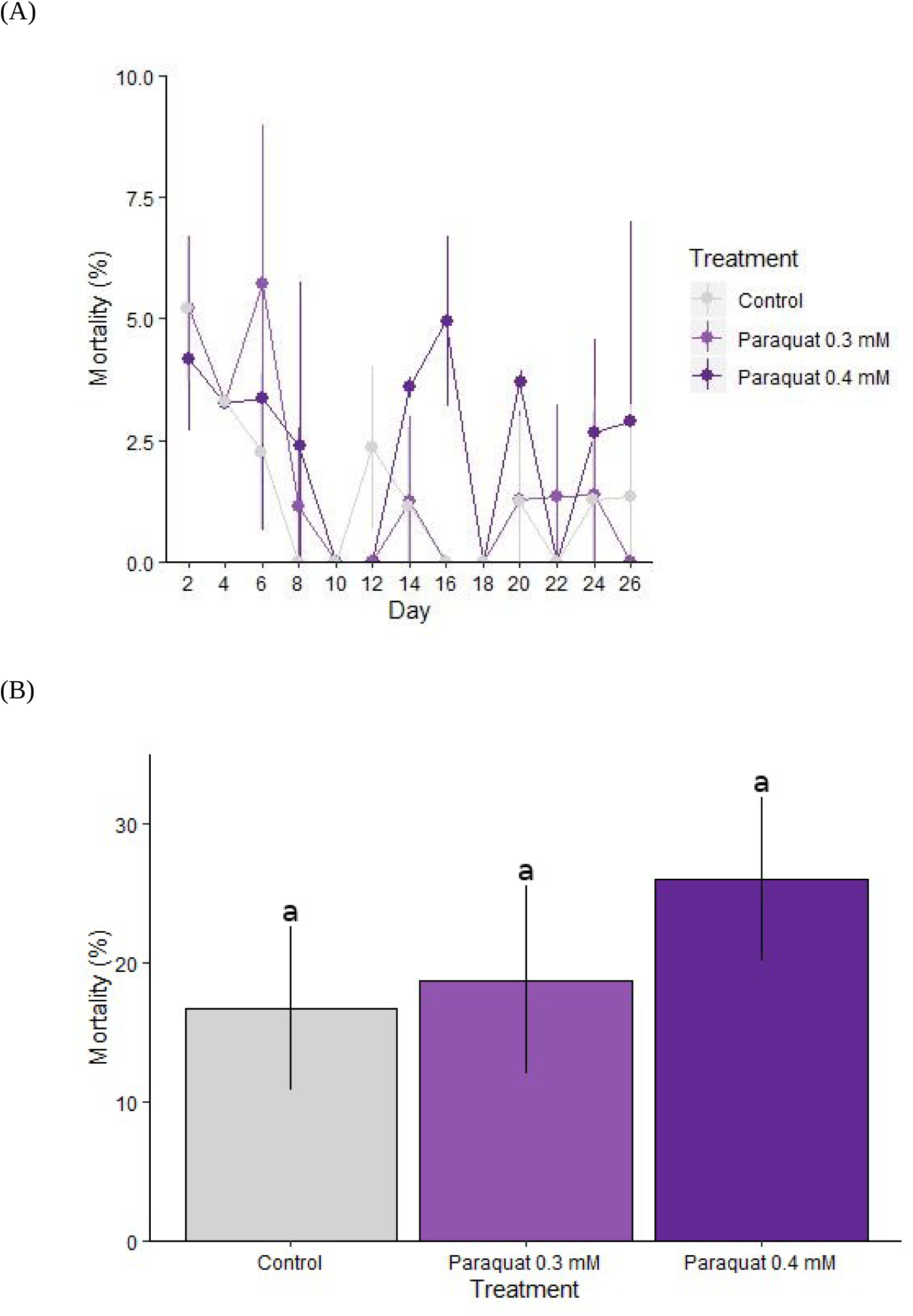
Mortality following stress exposure in *Helicoverpa armigera*. Individual *H. armigera* were reared on an artificial diet mixed with 0.3 mM or 0.4 mM paraquat and mortality was recorded from day two until 90% of the caterpillars had pupated. Results are presented as percentage mortality: (A) every two days; and (B) overall as a caterpillar, for each treatment and control group. Differences between treatment and control groups in (B) were not found to be statistically significant (i.e., all fall into a single homogeneous group). In both graphs, error bars indicate standard deviation and colours represent control or treated samples according to the provided key. Note the different *y-*axis scales across the two panels.

### Gene expression and ontology

#### Differential gene expression

An MDS plot of gene expression patterns among the top 1,000 most highly-expressed genes for 4th-instar caterpillars (both control and paraquat stressed samples) indicated that control and paraquat stressed samples show different patterns of gene expression, while 0.3 mM and 0.4 mM stressed samples cluster very similarly (Supporting Information Fig. S3). Therefore, we chose to analyse the data as stressed (n = 8) vs. control (n = 4) samples, although we also analysed pairwise comparisons (control vs. paraquat 0.3 mM and control vs. paraquat 0.4 mM) and obtained very similar results (data not shown).

In total, there were 1,618 up-regulated genes, 10,572 that were not significantly different, and 761 genes that were down-regulated in the DGE analysis. The full list of significantly (adjusted *P*<0.05) up- and down-regulated genes between control and stressed samples is shown in Supporting Information Table S2. Among the up-regulated genes were *caspase-4* and *meiosis arrest female 1 homolog protein*, while *collagen alpha-1(II) chain-like* gene was among the down-regulated (Supporting Information Table S2).

#### Gene ontology

Gene ontology (GO) analysis has the overall goal of identifying the functions of genes that are up- and down-regulated across treatment groups which are over-represented in the data. Functions are classified in terms of molecular function (MF), cellular component (CC), and biological process (BP). Overall, GO terms for 13 MF, two CC, and ten BP were enriched in the up-regulated gene set for paraquat stressed samples vs. controls, while GO terms for three BPs were enriched in the down-regulated gene set (Table 1).

**Table 1.**
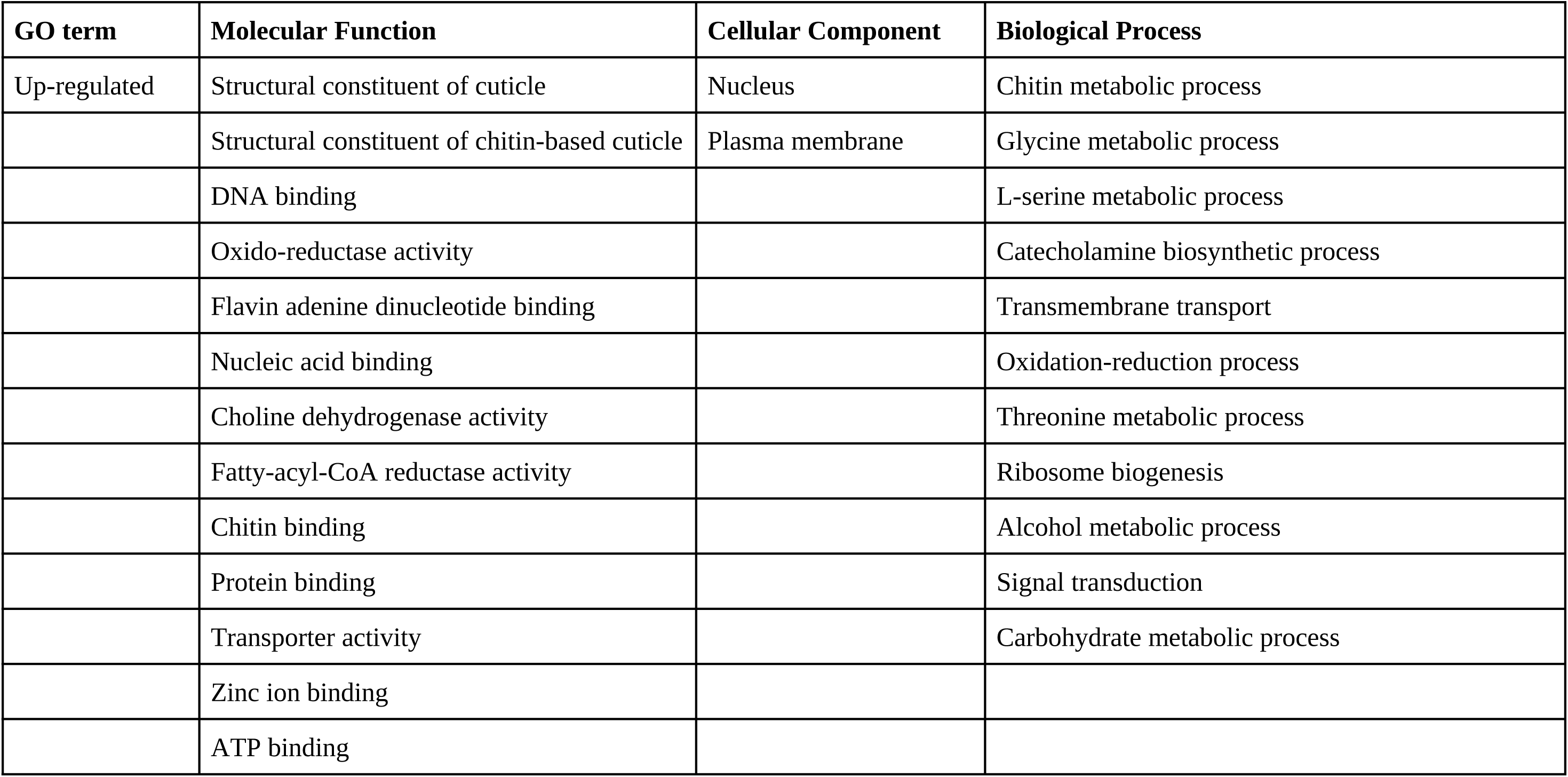
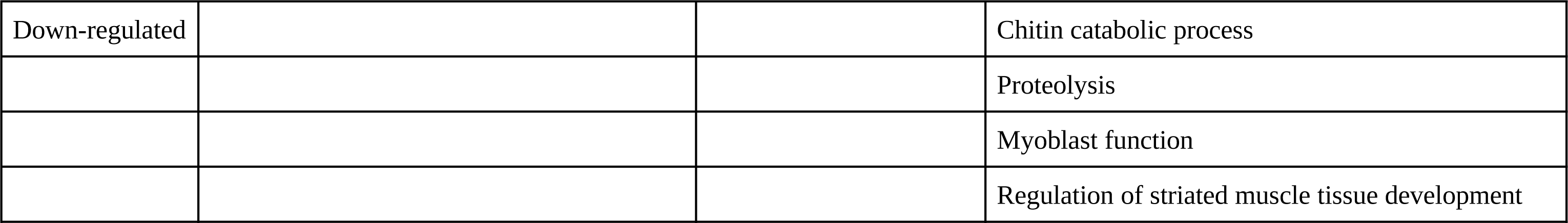
Significantly over represented gene ontology (GO) terms for up- and down-regulated genes in stressed vs. control samples of *Helicoverpa armigera*.

| GO term | Molecular Function | Cellular Component | Biological Process |
| --- | --- | --- | --- |
| Up-regulated | Structural constituent of cuticle | Nucleus | Chitin metabolic process |
|  | Structural constituent of chitin-based cuticle | Plasma membrane | Glycine metabolic process |
|  | DNA binding |  | L-serine metabolic process |
|  | Oxido-reductase activity |  | Catecholamine biosynthetic process |
|  | Flavin adenine dinucleotide binding |  | Transmembrane transport |
|  | Nucleic acid binding |  | Oxidation-reduction process |
|  | Choline dehydrogenase activity |  | Threonine metabolic process |
|  | Fatty-acyl-CoA reductase activity |  | Ribosome biogenesis |
|  | Chitin binding |  | Alcohol metabolic process |
|  | Protein binding |  | Signal transduction |
|  | Transporter activity |  | Carbohydrate metabolic process |
|  | Zinc ion binding |  |  |
|  | ATP binding |  |  |
| Down-regulated |  |  | Chitin catabolic process |
|  |  |  | Proteolysis |
|  |  |  | Myoblast function |
|  |  |  | Regulation of striated muscle tissue development |

For the MF subset, significantly (based on a False Discovery Rate; FDR>0.5) over-represented GO terms for up-regulated genes included oxido-reductase activity, chitin binding and cuticle, structural constituent of cuticle, flavin adenine dinucleotide binding, and fatty-acyl-CoA reductase (Table 1). The two over-represented GO terms corresponding to CC had functions related to nucleus and plasma membranes. Finally, the BP-related GO terms for the up-regulated gene set had functions involving glycine metabolism, transmembrane transport, ribosome biogenesis, signal transduction, and carbohydrate metabolism, and for the down-regulated gene set, were related to functions including proteolysis, regulation of striated muscle tissue development, and chitin catabolism (Table 1).

## Discussion

In this work, we investigated the physical and biochemical effects of paraquat-induced chronic oxidative stress on the pest moth *H. armigera*. We showed that exposure to moderate doses of paraquat slows development, elongates lifespan, and leads to the up-regulation of genes involved in detoxification (glycine metabolism), cuticle metabolism, and oxidation-reduction processes. However, no effect was observed on overall mortality, body weight, or time spent in other developmental stages.

### Developmental delay and extended lifespan

Our results suggest that chronic oxidative stress lengthens lifespan in *H. armigera* by slowing down development at the larval stages (∼25-28 days as larvae vs. ∼15 days, for stressed vs. control groups, respectively). Another study similarly showed that oxidative stress (paraquat) can slow pupal development and therefore lengthen lifespan in *H. armigera* (Zhang et al., 2017). Meanwhile, Pearce et al. (2017) found that lower quality host plants lead to slowed development rates in *H. armigera*. For example, a diet of *Arabidopsis* resulted in larvae requiring almost 13 days to reach 4th-instar, while larvae fed on cotton took only ∼8 days to reach the same developmental stage (Pearce et al., 2017). In addition, these authors found that, at 4th-instar, larvae fed on *Arabidopsis* weighed only ∼25 mg while larvae fed on a lab-diet weighed up to 50 mg. Combined with our results, this collective work suggests that developmental delay leading to lifespan extension under stress may be a common phenomenon in invertebrate species.

Indeed, the scenario of ROS extending lifespan has also been found in other invertebrates, including *D. melanogaster* (Hercus et al., 2003) and *C. elegans* (Lithgow & Walker, 2002). In the former study, the intriguing hypothesis of hormesis was raised to explain the fact that a compound that has an inhibitory or toxic effect at high doses can actually be beneficial at low doses (*sensu* Mattson, 2008). In fact, there is substantial research regarding hormesis due to the impacts of environmental stress on aging and longevity in invertebrate species (Le Bourg, 2009; Hunt et al., 2011; Scharf et al., 2017; Gilad et al., 2018; Mir & Qamar, 2018), including *Helicoverpa* (Ahn et al., 2011; Celorio-Mancera et al., 2011; Gulzar & Wright, 2015). In our case chronic oxidative stress extended the larval period in *H. armigera*, which is the developmental stage that causes the most damage to crops. This finding would therefore potentially have significant agricultural impacts should it occur in the wild, affecting the efficiency of reproduction in *H. armigera* (i.e., an extended caterpillar period would presumably delay reproductive events and lengthen the duration of vulnerability as larvae to natural enemies), and survival (i.e., prolonging exposure to predators, parasites, and disease). Further research (e.g. modelling of population growth under different times spent as a caterpillar) is needed in order to better determine the agricultural and potential economic effects of extended development in *H. armigera* under stressful conditions.

### Genes and pathways

Our results suggest that oxidative stress leads to retardation of developmental processes. The majority of research suggests that stress-induced cell death in *H. armigera* and other species occurs via the FoxO (Zhang et al., 2017) or p53 (Liu & Xu, 2011; Hori et al., 2013) pathways. However, we did not find genes in these pathways to be significantly differentially expressed. Instead, significant up-regulation of caspase and meiosis arrest female 1 genes was detected in response to stress and these genes were also differentially expressed across one or more diets when the larvae were raised on various stressful hosts in the work of Pearce et al. (2017). Caspases are a family of proteases that play a crucial role in inflammation and cell death (Galluzzi et al., 2016). Although the function of *caspase-4* is not yet fully understood, it may be associated with stress-induced apoptosis in Lepidoptera (Courtiade et al., 2011), and up-regulation of caspase proteins in response to stress has been shown in the diamondback moth, *Plutella xylostella* (Zhuang et al., 2011) and the Egyptian cotton leafworm, *Spodoptera littoralis* (Liu et al., 2005). Research on *meiosis arrest female 1* (*MARF1*) is limited - it is a novel vertebrate gene expressed exclusively in germ cells of the embryonic ovary and the adult testis (Arango et al., 2013). Though it is suggested to have a role in cell proliferation arrest in mice (Arango et al., 2013), and its up-regulation leads to deceleration of germline development in humans (Su et al., 2012), its’ role in insects, and in somatic development, is completely unclear. However, both *caspase-4* and *MARF1* appear to be important in stress responses in *H. armigera*, according to both our gene expression analyses and those of Pearce et al. (2017). These findings thus provide a basis for future work investigating the mechanisms underlying delayed development in response to oxidative stress in *H. armigera*.

Our gene expression analysis also identified an up-regulation of glycine metabolism, which has been shown to mitigate stress effects in mammals (Alves et al., 2019). Glycine is a non-essential amino acid involved in cryoprotection, anti-inflammation, and detoxification, and is also a crucial precursor of glutathione (an antioxidant molecule; Pérez-Torres et al., 2016). Thus, up-regulation of the glycine metabolic pathway may reduce the impact of toxic ROS on *H. armigera* during oxidative stress. An up-regulation of detoxification processes was also found in Pearce et al. (2017) following exposure to stressful diets. In total, these authors found 1,882 differentially expressed genes, of which 185 were from detoxification or digestion-related families (Pearce et al., 2017).

We also found an over-representation of genes involved in the structural constituent of cuticle, and of chitin metabolism, along with an under-representation of chitin catabolic processes in response to stress in *H. armigera*. Chitin functions to support the cuticles of the epidermis and trachea in insects, as well as the membranes that line the gut (Merzendorfer & Zimoch, 2003). Growth and cuticle-related genes also featured heavily in the up- and down-regulated gene lists of Pearce et al. (2017) and tens of these overlapped with those in our significant gene list. In general, 499 of the 1,882 differentially expressed genes in Pearce et al. (2017) overlapped with our set of 2,379, which is highly significant (hypergeometric test *P*=16.8 × 10^−22^) and suggestive of commonalities among transcriptomic stress responses, whatever the underlying trigger.

Interestingly, we also found that ecdysteroid (molting hormone) was up-regulated in stressed samples (1.5 fold change; *P=*0.002), while juvenile hormone (JH) was down-regulated (4.8 fold change; *P*=0.0009). Nutrition regulates growth and development in the majority of insects via levels of JH (Gotoh et al., 2014; Breed & Moore, 2015). Thus, lower nutrient absorption in stressed samples could be responsible for the down-regulation of JH if paraquat has damaging effects on the midgut (see Ahmad, 1995). Indeed, paraquat has known lipid peroxidation outcomes in invertebrates thus, lipid-dependent processes in insects are likely to be critically affected by oxidative stress - this includes the synthesis of ecdysone, JH, and other lipids that act as pheromones (Downer, 1985; Ahmad, 1995). Stress has been shown to interrupt hormone systems, resulting in down-regulation of JH in *Drosophila* (Kodrík et al., 2015), as well as honey bees (Lin et al., 2004) and the tobacco hawk moth, *Manduca sexta* (Tauchman et al., 2007). Depressed JH has also been shown to lead to a delay in ovary maturation in *Drosophila* (Saunders et al., 1990), and to longer diapause periods in flesh flies (Walker & Denlinger, 1980). In addition, recent research suggests that JH leads to increased levels of oxidative stress in the damselfly (Martínez-Lendech et al., 2019). Thus, molting and growth hormones could play a role in oxidative stress responses more generally and the relationship between hormone regulation and delayed development, as measured here in *H. armigera*, warrants further investigation.

Finally, a fundamental response to oxidative stress is the up-regulation of antioxidant enzymes, such as superoxide dismutase, catalase, and glutathione peroxidase, to reduce levels of oxidative damage by transforming ROS into non-toxic products (Halliwell, 1999; Mittler, 2002). Such responses are seen in an array of species (e.g., the southern armyworm, *Spodoptera eridania*, War et al., 2012; *D. melanogaster*, Arking et al., 2000), including *H. armigera* following exposure to high levels of UV-induced oxidative stress over time (Meng et al., 2009; Wang et al., 2012). However, though we did find an over-representation of genes involved in oxidation-reduction processes and nominal up-regulation, we did not find significant (*P*<0.05) up-regulation of these common antioxidant enzymes (superoxide dismutase: 3.1 fold change, *P*=0.075; glutathione peroxidase: 2.1 fold change; *P*=0.045).

### Summary

Previous research indicated that oxidative stress can impact cell senescence, apoptosis, and biochemical and metabolic pathways to have potentially strong effects on fitness. Here, we found that oxidative stress had marked effects on both development and gene expression in *H. armigera*. In particular, we found that, potentially linked to the hormesis hypothesis, sub-lethal paraquat exposure slowed down developmental rate, leading to a longer time spent as caterpillars and overall lifespan extension. Unresolved questions include whether this would be an advantage in the field and whether reproduction or other fitness-based traits were affected. At a molecular level, we further found that genes related to various developmental and detoxification processes were differentially expressed in response to stress. Collectively, these results advance our understanding of how *H. armigera* copes with stress and may help explain why this moth has become such a major pest.

## Acknowledgements

We would like to thank Megan Head for early advice on experimental design, NBA’s advisory panel at the Australian National University (Rod Peakall, Maja Adamska, Benjamin Schwessinger) for feedback on the early experimental design and the final thesis document, and Marcin Adamski and Darren Wong for assistance with gene expression, and gene ontology, analysis, respectively. We thank Chris Coppin for RNA library prep assistance, Amanda Padovan, Rachael Remington, and Theodore Colls for providing feedback on earlier drafts of this work, and Stephen Pearce and members of the Moritz lab group for helpful discussion. This project was supported through funding from the Australian Research Council (Discovery Early Career Researcher Award DE160100685 to AM), the Centre for Biodiversity Analysis (Ignition Grant to AM), and Commonwealth Scientific and Industrial Research Organisation Land and Water.

## Data Accessibility

Developmental data will be submitted to Dryad.

RNASeq data will be submitted to the Short Read Archive (SRA).

## Author Contributions

AM designed the research with input from NBA and TKW, NBA performed the developmental assays with assistance from BJ and created the libraries for RNASeq, NBA analysed the phenotypic data and AM analysed the RNASeq data, NBA compiled this work into her Master’s thesis with assistance from AM and TKW. AM re-worked and expanded the thesis text and figures into a manuscript and all authors contributed to the final version of the manuscript.

