## Supplementary Material for "Oxidative stress delays development and alters gene function in the agricultural pest moth, *Helicoverpa armigera*"

### Supporting Information

**Figure S1.** Mean weight following stress exposure in *Helicoverpa armigera*. Individual *H. armigera* were reared on an artificial diet mixed with 0.3 mM or 0.4 mM paraquat. Ten randomly-selected individuals were weighed every two days as caterpillars and twice a week as moths. Results are presented for stressed and control samples as mean weight every four days from day four. Differences between treatment and control groups were statistically significant - control samples had the highest mean weight from day four to day 16 ($F_{2,27}$=47.613, $F_{2,26}$=76.771, $F_{2,22}$=497.884, $F_{2,23}$=69.012, for days 4, 8, 12, and 16, respectively; $P$<0.001 for all) and from days 32 to 36 (day 32: $F_{2,17}$=34.874, day 36: $F_{2,17}$=14.297; $P$<0.001 for both) - but not between the two paraquat stressed groups, except for on day 20, when the 0.3 mM stressed samples were, on average, significantly heavier than their 0.4 mM counterparts. In the plot, significant differences among treatment and control groups are indicated by non-overlapping characters ('a', 'b'), error bars indicate standard deviation and colours represent control or treated samples according to the provided key.

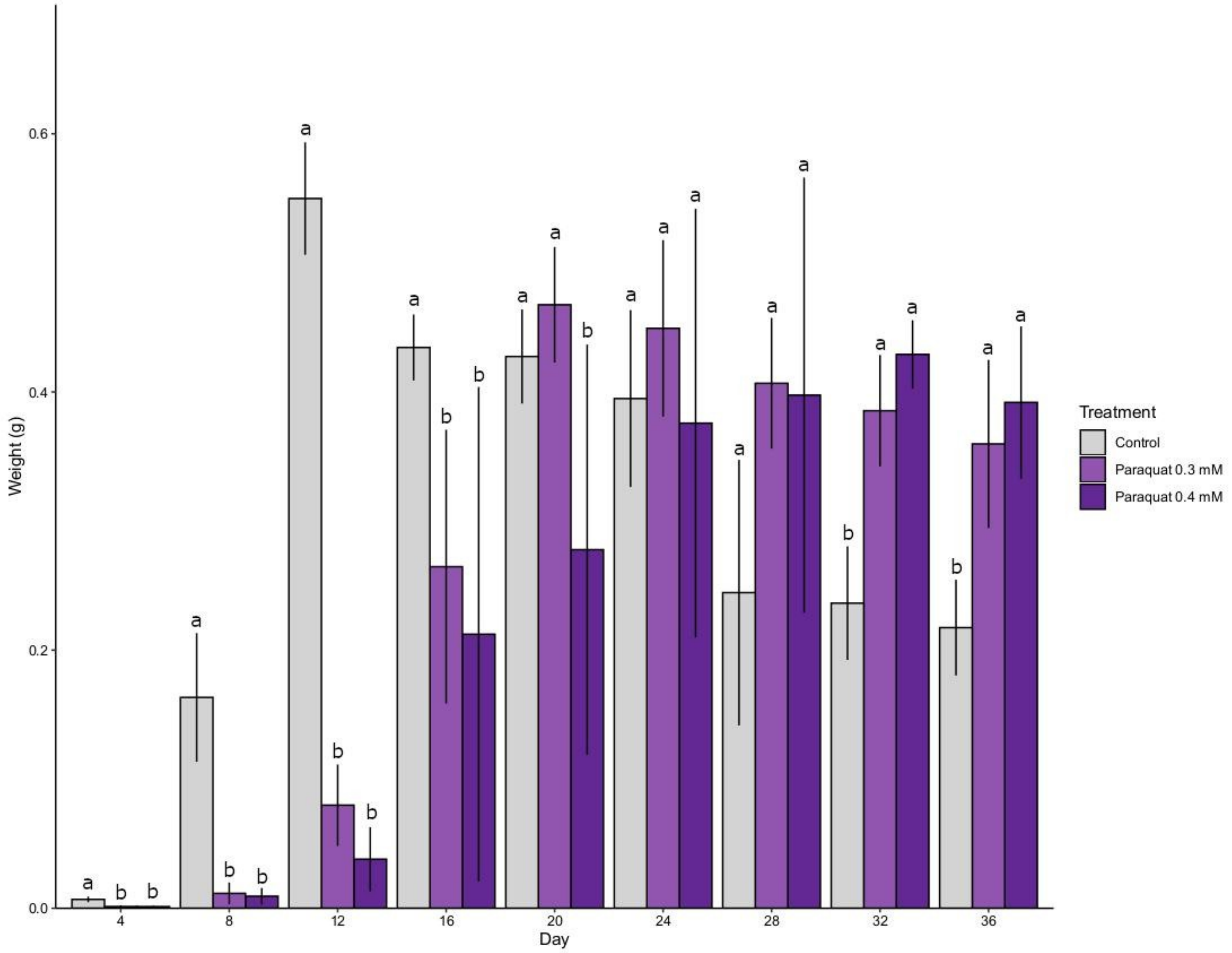

**Figure S2.** Developmental stage progression following stress exposure in *Helicoverpa armigera*. Individual *H. armigera* were reared on an artificial diet mixed with 0.3 mM or 0.4 mM paraquat and developmental stage was recorded every two days for control and stressed samples from the day that individuals hatched until death. Results are presented as the percentage of individuals representing each developmental stage (1st - 5th corresponding to instars, p-pu=pre-putation; pu=pupae) at: (A) day 12; and (B) day 20. Significant differences among treatment and control groups are indicated by non-overlapping characters (‘a’, ‘b’, ‘c’), error bars indicate standard deviation, and colours represent control or treated groups according to the provided key.

(A)

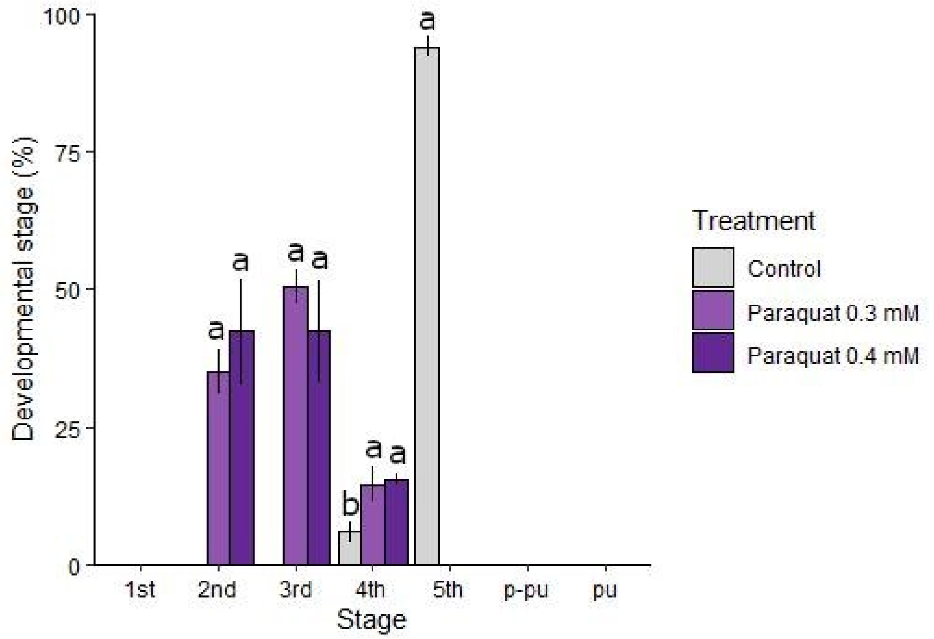


(B)

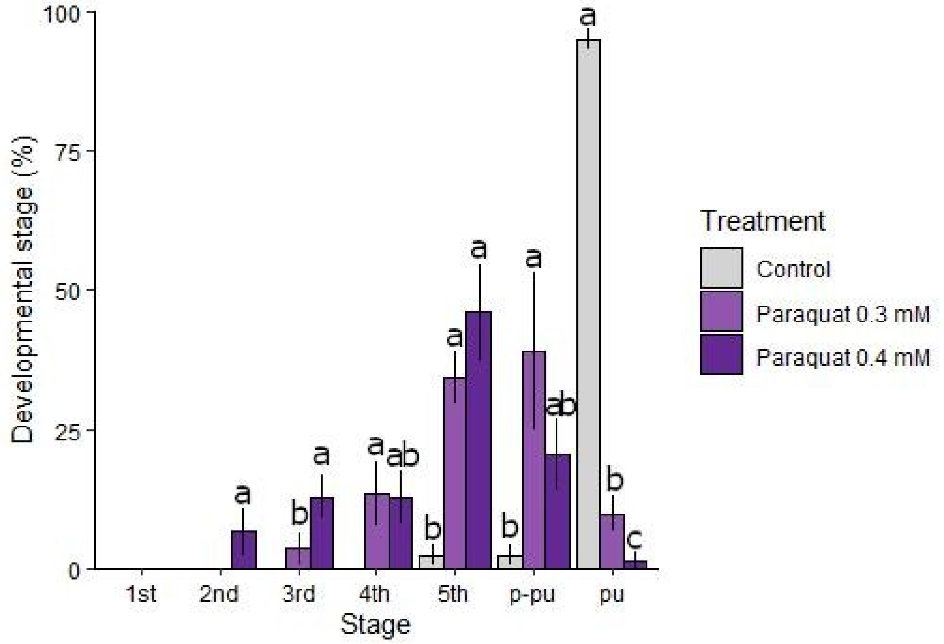

**Figure S3.** A multi-dimensional scaling plot of the top 1,000 differentially expressed genes; distances in the plot approximate the typical log2 fold changes between the samples, which consist of either controls (“C”) or 0.3 mM or 0.4 mM paraquat-exposed samples (“PQ3”, “PQ4”), respectively, as indicated by the provided colour key.

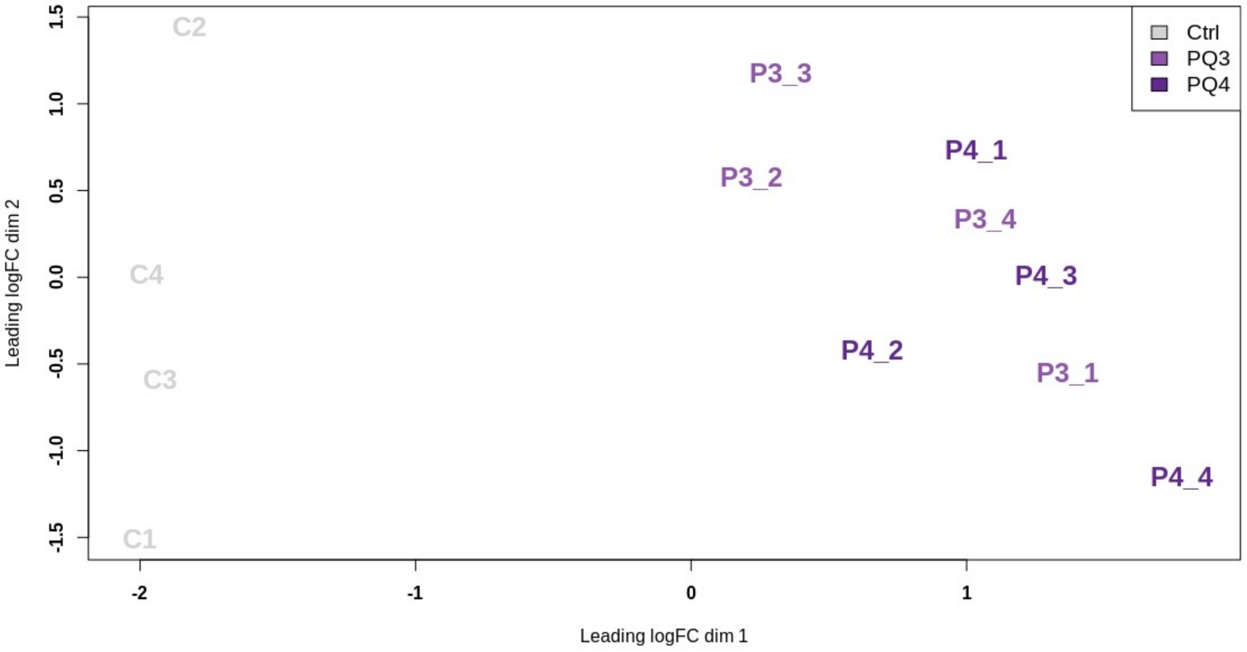

**Table S1.** Primer sequences.

| Primers | Sequences |
|---|---|
| RT_Hex[a] | CAGACGTGTGCTCTTCCGATCTNNNNNN |
| Bio_TS_RNA | Biotin-CAGGACGCTGTTCCGTTCAATggg[b] |
| TS_qPCR | CAGGACGCTGTTCCGTTCAATGGG |
| RT_Hex_qPCR[a] | CAGACGTGTGCTCTTCCGATCT |
| **Barcode primers**[c] | |
| i5_3 | AATGATACGGCGACCACCGAGATCTACACACAACCTCACACTTCGCTACAGGACGCTGTTCCGTTCAATGGG |
| i5_9 | AATGATACGGCGACCACCGAGATCTACACCTGTGAACACACTTCGCTACAGGACGCTGTTCCGTTCAATGGG |
| i5_11 | AATGATACGGCGACCACCGAGATCTACACGCGTATTCACACTTCGCTACAGGACGCTGTTCCGTTCAATGGG |
| i5_12 | AATGATACGGCGACCACCGAGATCTACACCGCATAAGACACTTCGCTACAGGACGCTGTTCCGTTCAATGGG |
| i7_A | CAAGCAGAAGACGGCATACGAGATCGAATTGCGTGACTGGAGTTCAGACGTGTGCTCTTCCGATCT |
| i7_B | CAAGCAGAAGACGGCATACGAGATGCTTAACGGTGACTGGAGTTCAGACGTGTGCTCTTCCGATCT |
| i7_C | CAAGCAGAAGACGGCATACGAGATCACCAAGAGTGACTGGAGTTCAGACGTGTGCTCTTCCGATCT |
| i7_E | CAAGCAGAAGACGGCATACGAGATGTGGTTCTGTGACTGGAGTTCAGACGTGTGCTCTTCCGATCT |
| i7_F | CAAGCAGAAGACGGCATACGAGATGGAGAGAAGTGACTGGAGTTCAGACGTGTGCTCTTCCGATCT |
| i7_G | CAAGCAGAAGACGGCATACGAGATCCTCTCTTGTGACTGGAGTTCAGACGTGTGCTCTTCCGATCT |
| i7_H | CAAGCAGAAGACGGCATACGAGATTTGTAGCCGTGACTGGAGTTCAGACGTGTGCTCTTCCGATCT |
| Custom Read 1 | ACACTTCGCTACAGGACGCTGTTCCGTTCAATGGG |

[a] Langevin, S. A., Bent, Z. W., Solberg, O. D., Curtis, D. J., Lane, P. D., Williams, K. P, Schoeniger, … Branda, S. S. (2013). Peregrine: a rapid and unbiased method to produce strand-specific RNA-Seq libraries from small quantities of starting material. *RNA Biology*, *10*, 502-515.
[b] Ribonucleotides are indicated with a lowercase letter.
[c] Barcode sequences are coloured.
